# Developmental benzo[a]pyrene exposure alters stress hormones, neurotransmitters and behavioral responses of mice dependent on *Cyp1* genotype

**DOI:** 10.1101/2025.08.13.670196

**Authors:** Jade Perry, Taylor Easybuck, Mackenzie Feltner, Emma G. Foster, Mickayla Kowalski, Amanda Honaker, Katelyn M. Clough, Alexandria Easton, Kevin Berling, Duong Pham, Annika White, Kayla Wypasek, Christine Perdan Curran

## Abstract

Benzo[a]pyrene (BaP) is a prototypical polycyclic aromatic hydrocarbon (PAH) produced during combustion processes and when grilling foods. Epidemiological studies indicate exposure to PAHs during pregnancy lead to learning and memory deficits as well as behavioral problems that persist into adolescence. Studies in rodents and zebrafish have frequently reported anxiolytic effects of BaP exposure in adult animals and in developmental studies. We conducted sequential experiments of *Cyp1a1(−/−)* and *Cyp1b1(−/−)* knockout mice compared with wild type C57BL/6J mice to determine if genotype changes the response to developmental BaP exposure. We treated pregnant dams from gestational day 10 to postnatal day 25 (P25) with BaP in corn oil-soaked cereal or the corn oil vehicle and tested one male and one female offspring beginning at P60. We found increased exploratory behavior in the elevated zero maze for *Cyp1a1(−/−)* knockout mice, but no significant differences in *Cyp1b1(−/−)* knockouts. In contrast, *Cyp1b1(−/−)* knockout mice buried fewer marbles in a second test of anxiety-like behavior. There were no significant differences when *Cyp1a1(−/−)* knockout mice were tested. BaP decreased immobility time in *Cyp1a1(−/−)* knockouts in the forced swim test, but increased immobility time in wild type and *Cyp1b1(−/−)* knockout mice. We measured plasma corticosterone levels at baseline and following the forced swim test and monoamine neurotransmitters at the end of behavioral testing. BaP treatment increased corticosterone in wild type mice, but decreased it in *Cyp1a1(−/−)* knockout mice. Both BaP-exposed and corn oil control *Cyp1b1(−/−)* knockout mice had higher corticosterone levels compared with wild type mice. Dopamine and serotonin signaling were altered in the hypothalamus dependent on genotype, treatment and sex. Together, these data suggest that both CYP1A1 and CYP1B1 have a normal role in brain functioning or development, and that CYP1 genotype alters the response to developmental BaP exposure in behavioral and biochemical tests related to stress, anxiety and depression.

**Highlights:**

- BaP-exposed *Cyp1a1(−/−)* mice had lower corticosterone and decreased immobility in the forced swim test
- BaP exposure increased FST immobility in wild type and *Cyp1b1(−/−)* mice
- *Cyp1a1(−/−)* and *Cyp1b1(−/−)* knockout mice showed less anxiety-like behavior
- Developmental BaP exposure altered corticosterone levels dependent on *Cyp1* genotype
- Genotype, treatment and sex all impacted neurotransmitter levels in the hypothalamus
- Genetic differences in CYP enzymes altered susceptibility to developmental BaP exposure

## 1. Introduction

Benzo[a]pyrene (BaP) is a widespread pollutant and known carcinogen produced from multiple combustion processes. BaP ranks in the top ten on the United States’ list of priority pollutants (ASTDR 2022). Human exposures are widespread from vehicle exhaust, fossil fuel-burning power plants, wildfire and tobacco smoke, and grilled food (ATSDR 1995). In recent years, evidence has been accumulating that developmental exposure to BaP and other polycyclic aromatic hydrocarbons (PAHs) can alter normal brain development with persistent, adverse effects during childhood and adolescence (Perera et al. 2012, 2018; Margolis et al 2016, 2021). In addition to lowered IQ at school age (Vishnevetsky et al. 2015), Perera et al. (2012) reported a positive association between prenatal PAH exposure and multiple behavioral outcomes including anxiety, depression and attention deficits in 6-7-year-old children. Those impairments and poor inhibitory control were recently associated with persistent problems in academic performance in adolescents (Margolis et al. 2021). In a meta-analysis of 16 epidemiological studies on human PAH exposure, Zhen et al. (2023) concluded that prenatal PAH exposure increased the risk of attention deficits nearly 3-fold and that PAH exposure increased the risk of depression in adults.

Numerous animal studies have confirmed that BaP exposure is strongly associated with not only learning and memory deficits, but persistent changes in behavior. Grova et al. (2008) used a range of doses from 0-200 mg/kg/day (i.p.) in adult female Balb/c mice for 10 days and found increased exploratory behavior in the hole board test and elevated plus maze at the 20 and 200 mg/kg/day dose. This study partially replicated their findings of an anxiolytic effect of high-dose BaP in young adult Balb/c female mice tested in the Y-maze (Grova et al. 2007). Similar anxiolytic effects were seen in female Swiss Albino mice dosed orally with 0.02 and 0.2 mg/kg/day BaP for 17 days (Bouayed et al. 2012). The BaP-treated mice showed similar behavior in the tail suspension test to the positive control animals treated with an anti-depressant and had higher levels of serotonin in the hippocampus and hypothalamus. Using a far different dosing approach, Das et al. (2017) directly injected 0.2 µg/kg of BaP into the cisterna magna of postnatal day 5 (P5) male Wistar rat pups prior to behavioral testing at P30. Similarly, these researchers reported anxiolytic effects with BaP-treated pups spending significantly more time in the open arms of the elevated plus maze and more time in the light zone in the light-dark text. Aparna et al. (2020) reported anxiolytic effects in adult zebrafish exposed to 0.2 mg/L of BaP with decreased time spent in the bottom during the novel tank diving test. Hamilton et al. (2021) found anxiolytic effects in zebrafish acutely dosed with 10 μM and 100 μM BaP. It’s important to note that none of these studies considered sex as a biological variable, which is a critical factor we addressed in our studies.

We previously demonstrated that both high-affinity *Ahr^b^Cyp1a2(−/−)* and poor-affinity *Ahr^d^Cyp1a2(−/−)* knockout mice were uniquely susceptible to developmental BaP neurotoxicity when tested as young adults (Honaker et al. 2022). Our earlier studies focused primarily on endpoints related to learning, memory and motor function. Our current studies were designed to determine if genetic differences exacerbate the neurotoxicity of developmental BaP exposure and its effects on measures of anxiety, depression and stress. All CYP1 enzymes have a potential role in BaP metabolism, but we focused these studies on *Cyp1a1(−/−)* and *Cyp1b1(−/−)* knockout mice based on earlier work demonstrating significant differences in BaP metabolism and immunotoxicity (Uno et al. 2004, 2006) and recent findings in humans that genetic differences in CYP1A1 increased the concentration of PAHs that crossed the placenta and accumulated in cord blood (Dong et al. 2018).

## 2. Methods

### 2.1 Animals

All animal experiments were conducted under protocols approved by the Northern Kentucky University Institutional Animal Care and Use Committee (IACUC) and follow the ARRIVE Guidelines (Percie du Sert et al. 2020; Kilkenny et al. 2010). C57BL/6J (wild type for all genotypes) mice were purchased from The Jackson Laboratory (Bar Harbor, ME), *Cyp1a1(−/−)* (Dalton et al. 2000) and *Cyp1b1(−/−)* (Buters et al. 1999) mouse lines were used for the current study. The knockout lines were routinely back-crossed to C57BL/6J mice to avoid confounding by genetic drift. Genotypes were confirmed through PCR genotyping from tail snips at weaning and after sacrifice.

#### 2.1.1 Animal husbandry

Animal housing and care followed the Guide for the Care and Use of Laboratory Animals (8th edition). All mice were maintained in the Northern Kentucky University vivarium. Mice were housed in polysulfone shoebox cages with corncob bedding and one cotton nestlet as enrichment. Up to 4 adult mice were housed together, grouped by sex, genotype, and treatment. Water and Lab Diet 5015 chow (18.9% protein, 11% fat, 32.6% starch) were provided *ad libitum* and lighting was set to a 12 h: 12 h light-dark cycle. Cages were changed weekly, and mice were checked daily for any possible health concerns.

#### 2.1.2 Breeding

Female and male mice of the same genotype were mated over a 4-day breeding cycle. The day a vaginal plug was found was considered gestational day 0.5 (G0.5), and the female mouse was removed from the breeding cage. To control for maternal care, litters were balanced at six pups per dam by either culling or cross-fostering pups of the same genotype and treatment group.

### 2.2 Benzo[a]pyrene treatments

Benzo[a]pyrene (> 96% purity; Millipore-Sigma) was dissolved in corn oil. Beginning at G10, pregnant dams were randomly assigned to treatment groups. The BaP group was given 10 mg/kg/day BaP dissolved in corn oil while the control group received an equivalent volume of the corn oil vehicle. To avoid the stress of daily injections or gavage, dams ingested the treatments on small pieces of cereal (Cap’n Crunch^TM^ peanut butter). Treatment of dams continued until P25 when the pups were weaned, so all exposures to the offspring were in utero and via lactation.

### 2.3 Behavioral testing

At P25, one male and one female pup from each litter were randomly selected for behavioral testing beginning at P60. Animals from all treatment groups were tested together in overlapping cohorts to avoid confounding by seasonal differences in behavior. Only one test was conducted per day within a 4 h time period to avoid confounding by circadian rhythms. Experiments were conducted under reduced lighting with no investigators present, and all investigators were blinded to treatment for animal handling and scoring. For all tests, 15-20 litters per group were used with a single trial for each animal. When multiple animals were tested simultaneously, only animals of the same sex were present in the room. Experiments are described in the order they were conducted. We tested and analyzed each knockout line separately; therefore, results are presented sequentially and separately.

#### 2.3.1 Elevated zero maze

As previously described (Brown-Villalona et al. 2020), a circular maze (105 cm diameter) with two open and two closed quadrants was used to assess anxiety-like behavior and hyperactivity. Mice were placed in a closed quadrant and behavior was recorded for 5 min. The videos were then scored using ODlog^TM^ for the following: latency to first leave the closed quadrant, head dips over the sides of the maze, zone crossings from one closed quadrant to another, and total time spent in the open quadrants.

#### 2.3.2 Marble burying

Mice were placed in a polysulfone shoebox cage filled with 5 cm of woodchip bedding and 15 identical marbles evenly placed across the bedding in a 3 x 5 grouping using a template. The mice were left alone for 20 min before being removed. The number of marbles buried at least 2/3 were recorded as buried, and pictures of each cage were taken to ensure accurate scoring.

#### 2.3.3 Forced swim test

Mice were placed in 10 cm opaque cylinders filled with water (25±1°C), and their behavior was recorded for 6 min. Videos were scored using ODLog ^TM^ for the following: latency to start floating and the total time spent floating, which was used to calculate the percent time immobile.

### 2.4 Corticosterone assays

Blood was collected from the saphenous vein of each animal at baseline and 5 min after the forced swim test, centrifuged, and the plasma stored at −80°C until analysis. We used an ELISA kit (Alpco Diagnostics) to quantify corticosterone levels following the vendor’s protocol.

### 2.5 Neurotransmitter analysis

Animals were sacrificed by decapitation following behavioral testing ∼P120. Brain regions were dissected, snap frozen and stored at −80°C until analysis. Methods were modified from Gutierrez et al. (2018). Tissues were weighed and homogenized in 25 volumes of 0.2N perchloric acid with a glass Dounce homogenizer and centrifuged. Monoamine neurotransmitters dopamine and serotonin and their metabolites, 3,4-dihydroxyphenylacetic acid (DOPAC) and 5-hydroxyindolacetic acid (5HIAA), were quantified using high-performance liquid chromatography (Waters Alliance e2695) with electrochemical detection (Waters 2465). Using an autosampler, 10 µl of supernatant was injected onto a Waters XBridge C18 column (3.5 µm, 4.6 x 150 mm) with isocratic MD-TM mobile phase (Thermo Fisher Scientific) and a constant flow rate of 0.5 mL/min. Using Empower software (Waters, Inc. Milford, MA), chromatograms were integrated and analyzed with neurotransmitter concentrations calculated based on standard calibration curves. Data from the hypothalamus analysis are presented here.

### 2.6 Statistical analysis

IBM SPSS statistical software (Version 28.0) was used to analyze all data for main effects of genotype, treatment and sex as well as their interactions. When differences were found, post-hoc analyses were done using corrections for multiple comparisons. We considered significance when P < 0.05. Data are presented as least square means ± the standard error of the mean (SEM). Behavioral tests included a minimum of 15 litters per group. Biochemical assays included a minimum of 8 litters per group. These group sizes were consistent with our prior experience using highly inbred mouse lines.

## 3. Results

### 3.1 Elevated zero maze

We found a main effect of genotype and a main effect of sex (F_1,184_ = 4.69, P < 0.05) in the elevated zero maze test. *Cyp1a1(−/−)* knockout mice spent significantly more time in the open quadrants (F_1,184_ = 24.12, P < 0.001) and completed significantly more zone crossings (F_1,184_ = 11.54, P < 0.01) compared with *Cyp1a1(+/+)* wild type mice (Figs. 1A-B). Female mice spent significantly more time in the open compared with male mice regardless of treatment or genotype (data not shown). We did not find any effects of genotype or treatment when testing *Cyp1b1(−/−)* knockout mice, but we did replicate our findings of sex differences with female mice spending more time in the open (F_1,12 0_ = 5.66, P < 0.05) and completing more zone crossings (F_1,120_ = 14.27, P < 0.001) compared with male mice (data not shown).

**Fig. 1.**
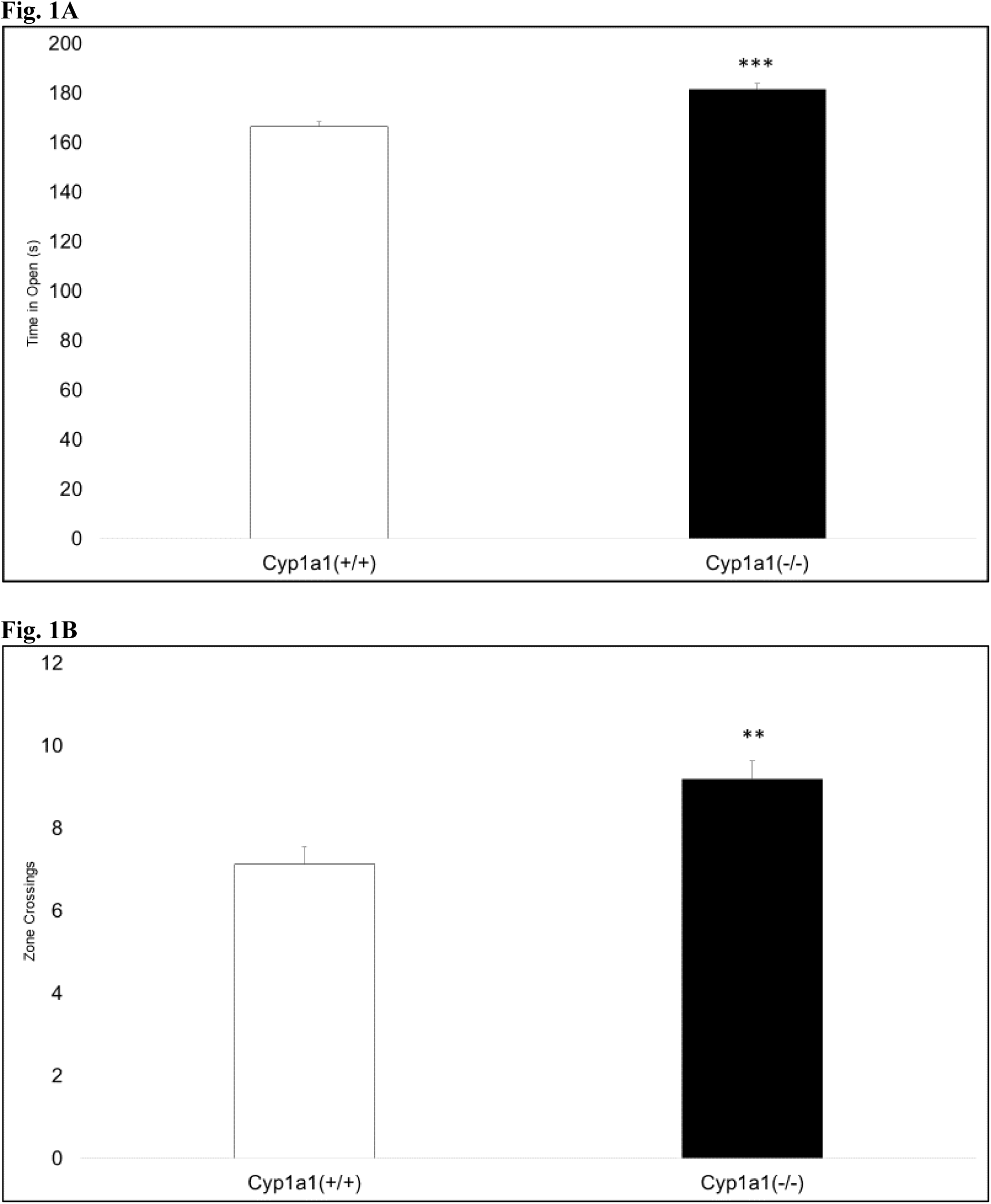
Elevated zero maze. There was a main effect of genotype with *Cyp1a1(−/−)* knockout mice spending more time in the open (A) and completing more zone crossings (B) compared with *Cyp1a1(+/+)* wild type mice. ** P < 0.01, *** P < 0.001

### 3.2 Marble burying

There were no significant main effects or interactions in the marble burying test (P > 0.05) when comparing BaP-exposed and control *Cyp1a1(−/−)* knockout and *Cyp1a1(+/+)* wild type mice (Fig. 2A). In contrast, we found that *Cyp1b1(−/−)* knockout mice buried significantly fewer marbles (F_1,120_ = 8.90, P < 0.01) than *Cyp1b1(+/+)* wild type mice (Fig. 2B). We note there were differences in the wild type (C57BL/6J) results across the two experiments. This might indicate a difference in how investigators interpreted what constitutes a 2/3 buried marble.

**Fig. 2.**
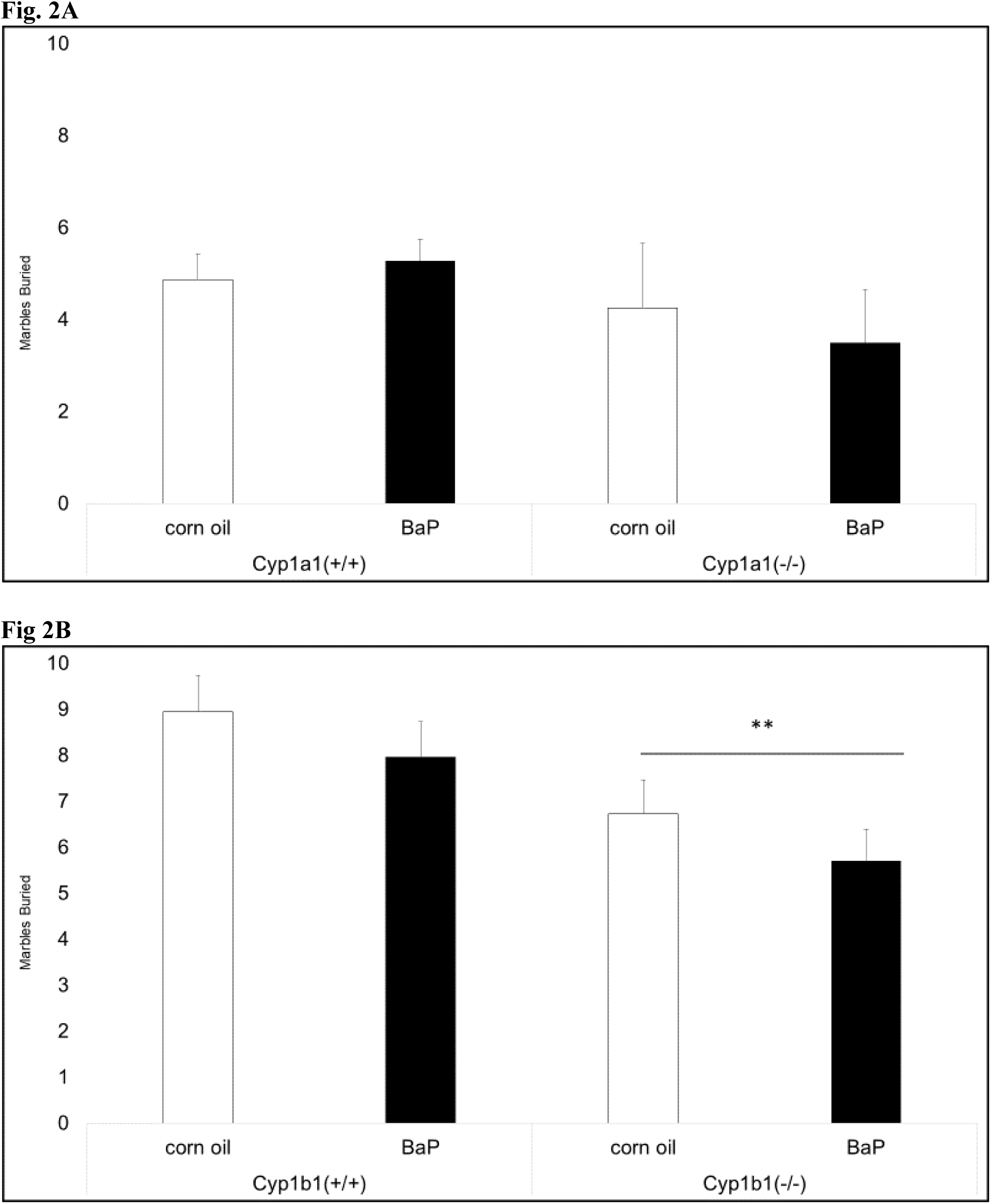
Marble burying. There were no significant differences in the first experiment comparing *Cyp1a1(−/−)* knockout and *Cyp1a1(+/+)* wild type mice (A). There was a main effect of genotype in the second experiment with *Cyp1b1(−/−)* knockout mice burying fewer marbles compared with wild type mice (B). ** P < 0.01

### 3.3 Forced swim test (FST)

We found a significant gene x treatment interaction in the forced swim test (F_1,140_ = 4.10, P < 0.05) when comparing *Cyp1a1(−/−)* knockout and *Cyp1a1(+/+)* wild type mice. BaP-exposed *Cyp1a1(−/−)* knockout mice spent less time immobile compared with their corn oil-treated controls whereas BaP-exposed *Cyp1a1(+/+)* mice floated more than their corn oil controls (Fig. 3A). We found significant main effects of genotype (F_1,120_ = 4.93, P < 0.05) and treatment (F_1,120_ = 5.82, P < 0.05) when comparing *Cyp1b1(−/−)* knockout and *Cyp1b1(+/+)* wild type mice. *Cyp1b1(−/−)* knockout mice spent significantly less time immobile compared with *Cyp1b1(+/+)* wild type mice, and BaP-exposed mice floated significantly more than corn oil-treated controls (Fig. 3B).

**Fig. 3.**
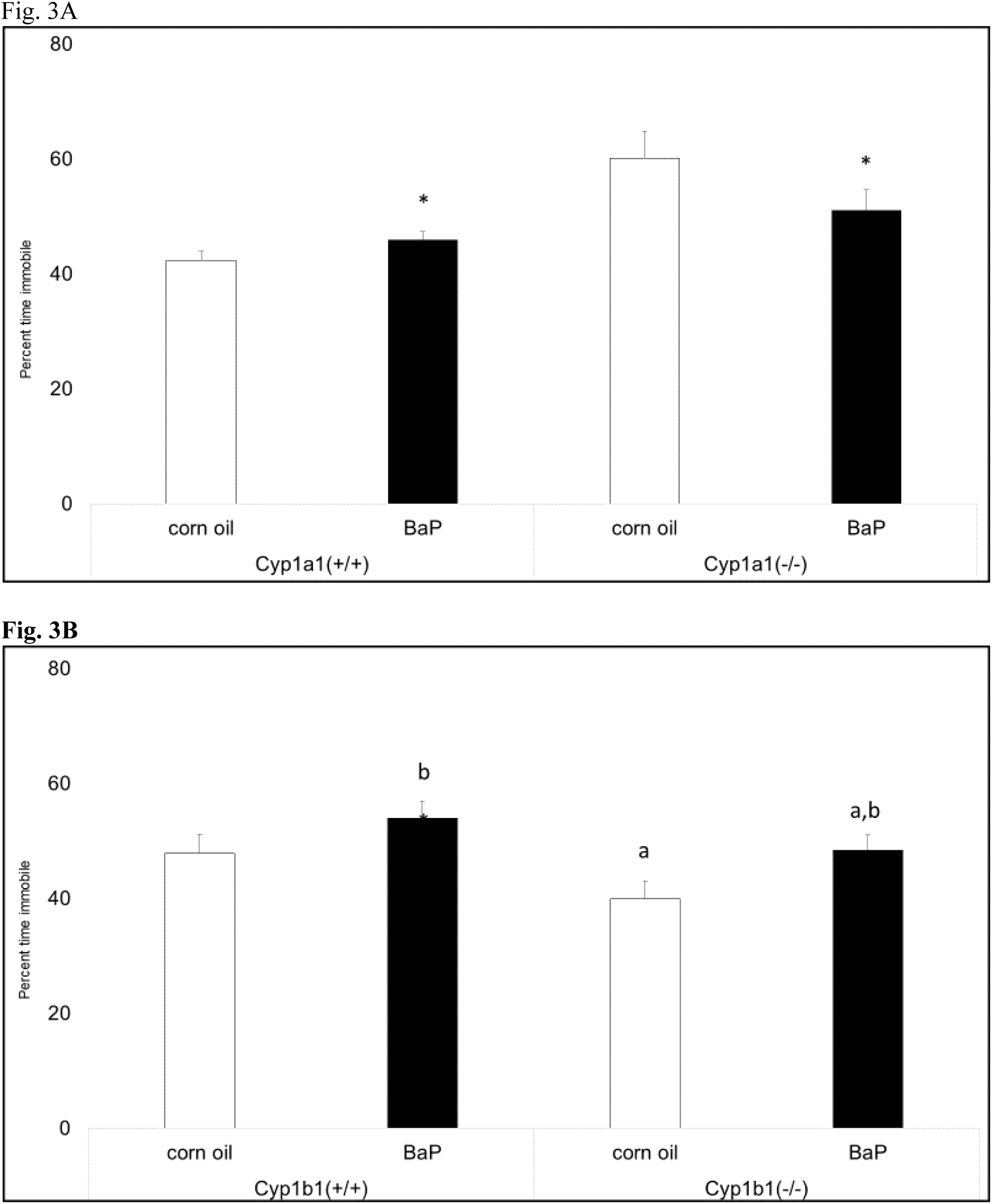
Forced swim test. There was a gene x treatment interaction in the first experiment with BaP-exposed *Cyp1a1(+/+)* wild type mice spending slightly more time immobile than their corn oil-treated controls whereas BaP-exposed *Cyp1a1(−/−)* knockout mice spent less time immobile compared with their corn oil-treated controls (A). In the second experiment, there was a main effect of genotype with *Cyp1b1(−/−)* knockout mice spending less time immobile than wild type mice (B). * P < 0.05, a = significantly different from wild type, b = significantly different from corn oil.

### 3.4 Plasma corticosterone levels

We found a significant gene x treatment interaction for baseline corticosterone levels (F_1,32_ = 4.25, P < 0.05) as well as for levels following the stress of the forced swim test (F_1,85_ = 4.79, P < 0.05) when comparing *Cyp1a1(−/−)* knockout and *Cyp1a1(+/+)* wild type mice. Baseline levels were significantly higher in BaP-exposed *Cyp1a1(+/+)* wild type mice, but lower in BaP-exposed *Cyp1a1(−/−)* knockout mice compared with corn oil controls (Fig. 4A). There was a similar trend in corticosterone levels following the stress of the forced swim test (Fig. 4B). In contrast, we found corticosterone levels were significantly higher in *Cyp1b1(−/−)* knockout mice (F_1,61_ = 15.78, P < 0.001) after the forced swim test regardless of treatment (Fig. 4C).

**Fig. 4.**
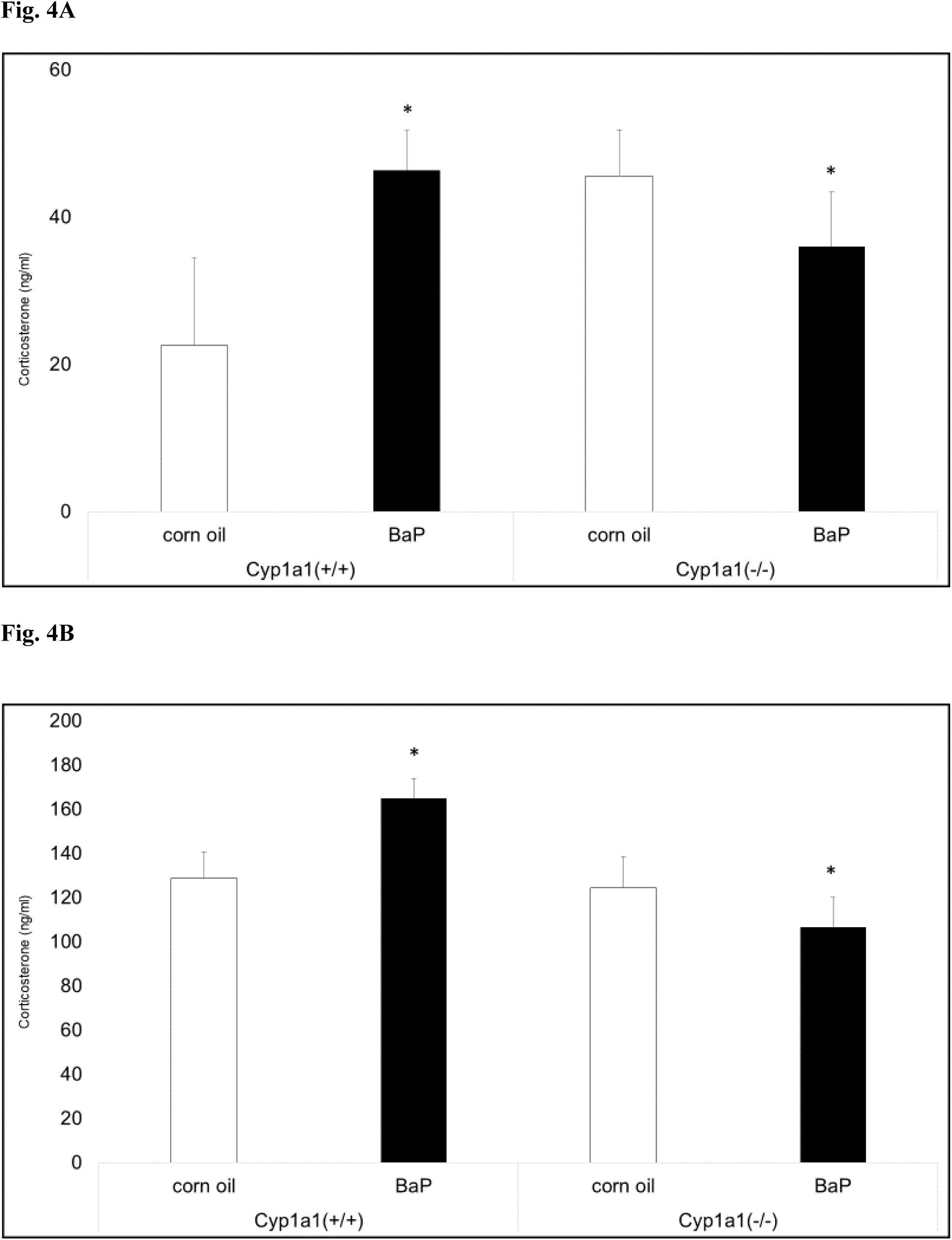
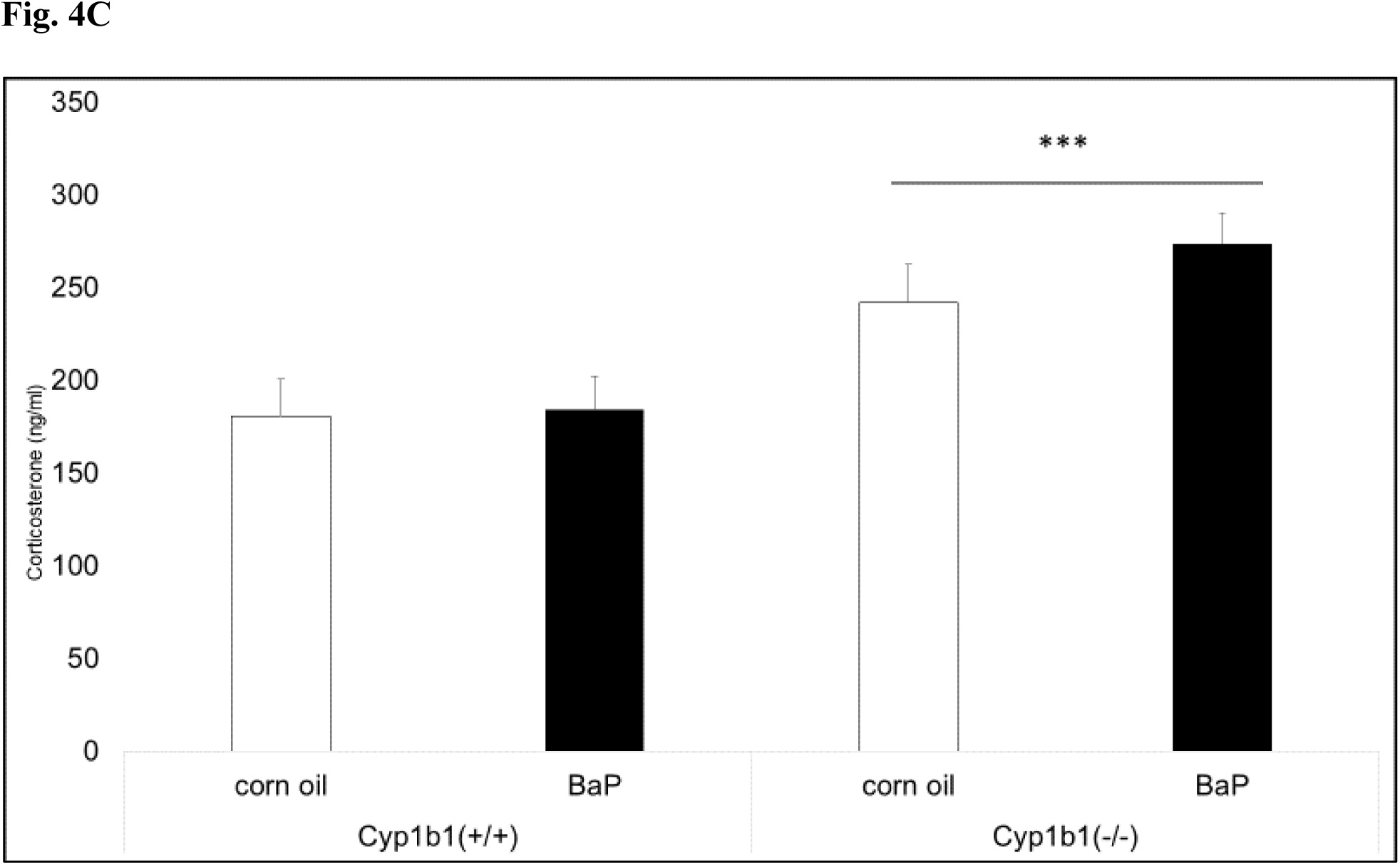
Corticosterone levels in plasma. There was a significant gene x treatment interaction in the first experiment with BaP-treated *Cyp1a1(−/−)* knockout mice having lower levels of corticosterone compared with their corn oil-treated controls while BaP-treated *Cyp1a1(+/+)* wild type mice had significantly higher levels than corn oil controls at both baseline (A) and following the forced swim test (B). There was a main effect of genotype in the second experiment with *Cyp1b1(−/−)* knockout mice having higher levels of corticosterone following the forced swim test (C). * P < 0.05, *** P < 0.001

### 3.5 Hypothalamus neurotransmitter levels

There was a significant gene x treatment x sex interaction for dopamine (F_1,181_ = 4.18, P < 0.05) and its metabolite DOPAC (F_1,81_ = 4.96, P < 0.05) in *Cyp1a1* knockout and wild type mice. Corn oil-treated *Cyp1a1(−/−)* female mice had significantly higher dopamine levels than all other groups, and BaP-exposed male *Cyp1a1(+/+)* wild type mice had significantly reduced dopamine compared with females. (Table 1). BaP-exposed male *Cyp1a1(+/+)* wild type mice and corn oil-treated *Cyp1a1(−/−)* male mice had significantly lower DOPAC levels compared with the corresponding females (Table 2).

**Table 1.**
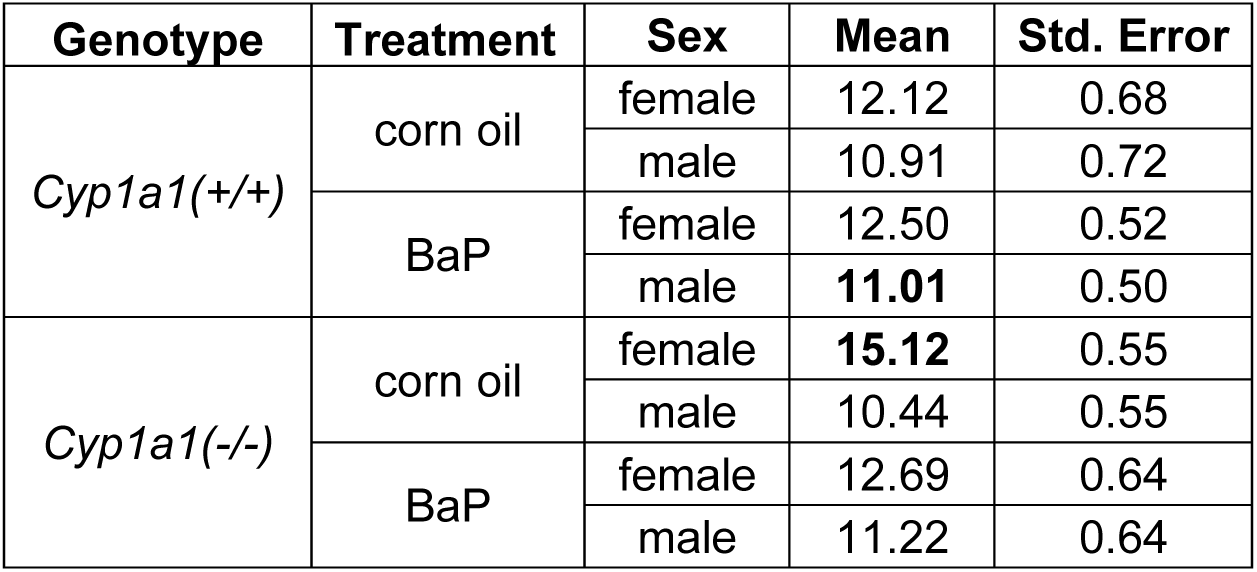
Dopamine levels in the hypothalamus.

| Genotype | Treatment | Sex | Mean | Std. Error |
| --- | --- | --- | --- | --- |
| *Cyp1a1*(+/+) | corn oil | female | 12.12 | 0.68 |
|  |  | male | 10.91 | 0.72 |
|  | BaP | female | 12.50 | 0.52 |
|  |  | male | **11.01** | 0.50 |
| *Cyp1a1*(-/-) | corn oil | female | **15.12** | 0.55 |
|  |  | male | 10.44 | 0.55 |
|  | BaP | female | 12.69 | 0.64 |
|  |  | male | 11.22 | 0.64 |

**Table 2.** DOPAC levels in hypothalamus.

| Genotype | Treatment | Sex | Mean | Std. Error |
| --- | --- | --- | --- | --- |
| *Cyp1a1*(+/+) | corn oil | female | 3.41 | 0.16 |
|  |  | male | 3.40 | 0.17 |
|  | BaP | female | 3.50 | 0.12 |
|  |  | male | **3.05** | 0.12 |
| *Cyp1a1*(-/-) | corn oil | female | 3.33 | 0.13 |
|  |  | male | **2.62** | 0.13 |
|  | BaP | female | 3.24 | 0.15 |
|  |  | male | 2.97 | 0.15 |

There were no significant differences in serotonin levels in the hypothalamus when comparing *Cyp1a1* knockout and wild type mice; however, there were significant main effects of genotype (F_1,81_ = 18.6, P < 0.001), treatment (F_1,81_ = 5.44, P < 0.05) and sex (F_1,81_ = 79.63, P < 0.001) for the serotonin metabolite 5-HIAA. *Cyp1a1(−/−)* knockout mice had significantly lower levels of 5-HIAA compared with *Cyp1a1(+/+)* wild type mice (Fig. 5A), and BaP-exposed mice had significantly lower levels of 5-HIAA compared with corn oil-treated mice (Fig. 5B). Female mice had significantly higher levels of 5-HIAA compared with male mice (data not shown).

**Fig. 5.**
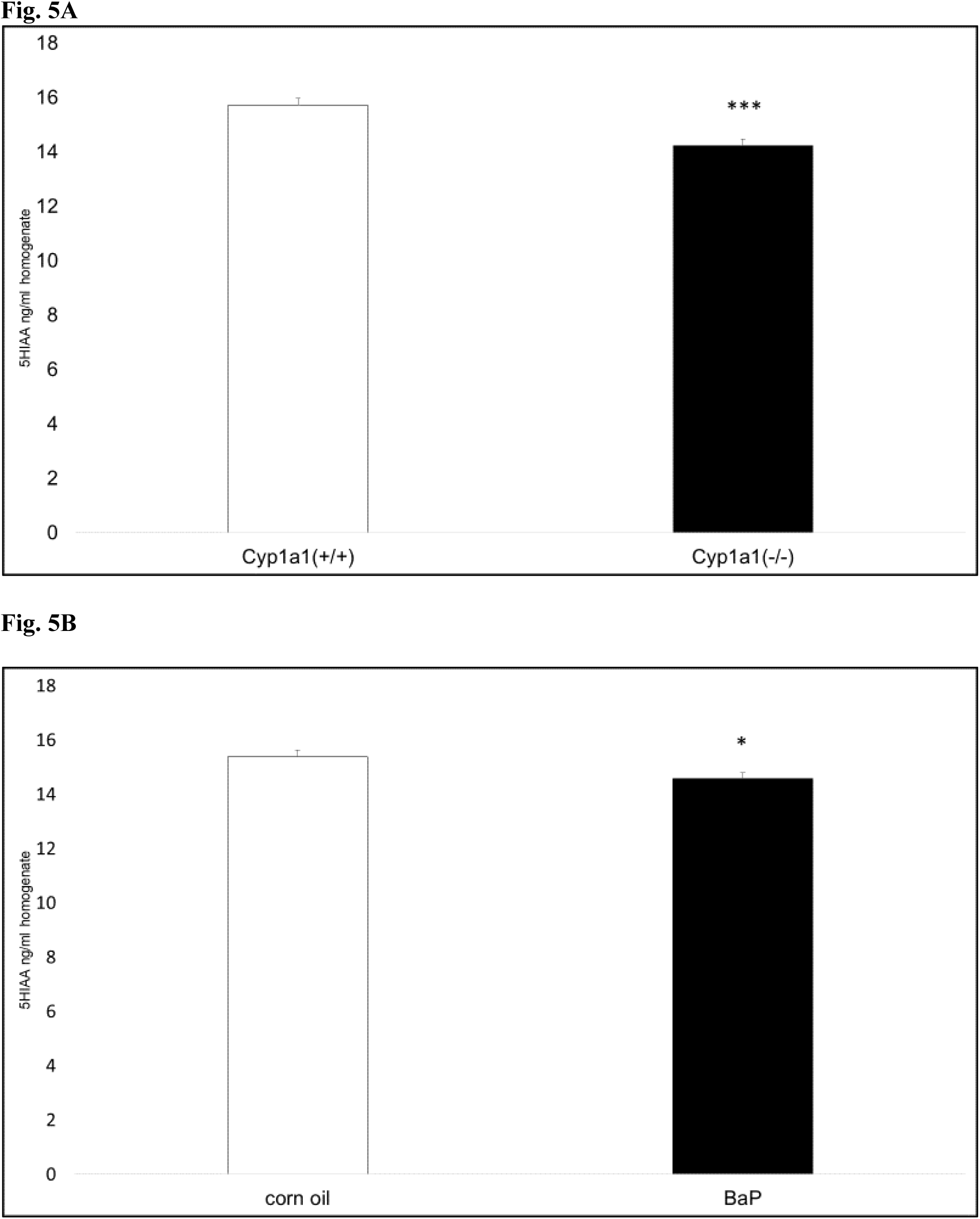
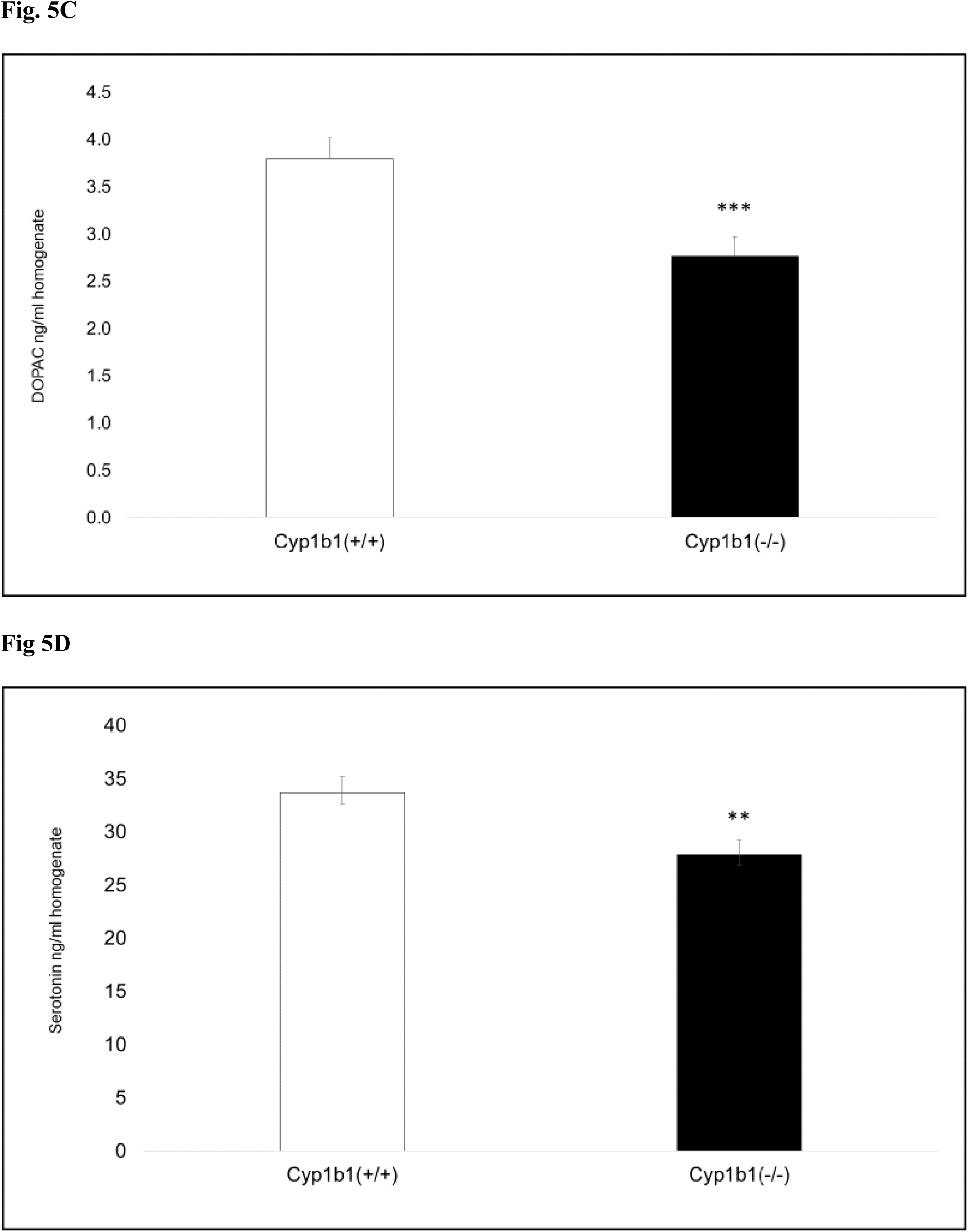
Monoamine neurotransmitters in the hypothalamus. There was a main effect of genotype in the first experiment with significantly lower levels of the serotonin metabolite 5-HIAA in *Cyp1a1(−/−)* knockout mice (A). BaP exposure significantly lowered 5-HIAA levels in the first experiment (B). In the second experiment, the dopamine metabolite DOPAC (C) and serotonin levels (D) were both significantly lower in *Cyp1b1(−/−)* knockout mice compared with wild type mice. * P < 0.05, ** P < 0.01, *** P < 0.001

Genotype was highly significant in the hypothalamus of *Cyp1b1(−/−)* mice with lower levels of DOPAC (F_1,119_ = 11.32, P < 0.01) (Fig. 5C) and a trend for significance for dopamine (F_1,119_ = 3.66, P =0.058). Serotonin levels were also significantly lower in *Cyp1b1(−/−)* mice (F_1,119_ = 8.15, P < 0.01) (Fig. 5D), but there was no difference in the levels of 5-HIAA (data not shown). There was no effect of BaP treatment for any neurotransmitter or metabolite in this second study.

## 4. Discussion

Prenatal exposure to polycyclic aromatic hydrocarbons (PAHs) has been linked to deficits in academic performance and behavior in highly exposed human populations (Perera et al. 2012, Vishnevetsky et al. 2015, Margolis et al 2016, 2021). We have completed sequential studies of *Cyp1a1(−/−)* and *Cyp1b1(−/−)* knockout mice exposed to the well-characterized PAH benzo[a]pyrene (BaP) during early brain development with behavioral testing conducted on young adults. Both knockout lines were back-crossed more than 8 generations onto a C57BL/6J (B6) background, and B6 mice were used as the wild type in all experiments. Multiple rodent studies indicated that BaP exposure in adult and neonatal animals produces an anxiolytic effect in commonly used tests of anxiety-like and depressive-like behavior (Grova et al. 2007, 2008; Bouayed et al. 2009; Das et al. 2017). The goals of our experiments were to determine if genetic differences altered the effects of developmental BaP exposure on adult behavior and to determine if males and females were differentially affected because prior studies did not include sex as a biological variable. We hypothesized that both knockout lines would be more susceptible due to reduced capacity to metabolize and clear BaP in the dams and pups.

We used complementary tests of anxiety-like behavior (elevated zero maze and marble burying), and both knockout lines showed less anxiety-like behavior than wild type mice regardless of BaP exposure. However, the results across tests were not consistent. *Cyp1a1(−/−)* knockout mice showed increased exploratory behavior in the zero maze (Fig. 1A-B), but no differences in the marble burying test (Fig. 2A). In contrast, *Cyp1b1(−/−)* mice were no different than wild type mice in the zero maze, but showed significantly less anxiety-like behavior in the marble burying test (Fig. 2B). We found no evidence of an anxiolytic effect of BaP, but we did find sex differences with females in both experiments demonstrating more exploratory behavior in the zero maze. Together, these findings suggest that the genotype and sex are more important than BaP treatment for outcomes related to anxiety-like behavior and that both CYP1 enzymes have some role in brain development or function.

The forced swim test (FST) is commonly used to test depressive-like behavior in rodents and the effectiveness of antidepressant medications (Colletis et al. 2024, Vahid-Ansari et al. 2024). We found different trends in the two knockout lines of mice with BaP-exposed *Cyp1a1(−/−)* knockout mice having decreased immobility time compared to their corn oil controls (Fig. 3A). *Cyp1b1(−/−)* mice spent significantly less time immobile than wild type mice (Fig. 3B).

Unexpectedly, we found that 10/mg/kg/day BaP treatment in dams increased immobility time in wild type and *Cyp1b1(−/−)* knockout offspring compared to their corn oil controls. This would be consistent with the findings of Zhang et al. (2016) who reported increased anxiety-like behavior in C57BL/6 adult male mice chronically dosed with 6.25 mg/kg/day BaP. In contrast, Bouayed et al. (2012) reported an anxiolytic effect of lower doses of BaP (0.02 and 0.2 mg/kg/day) in adult female Swiss albino mice in the tail suspension test, but no differences in the FST. This indicates that age, genetic background and dose all play a role in the behavioral outcomes.

To help uncover neuroendocrine signaling that might underlie the behavioral differences, we measured corticosterone levels at baseline and 5 min following the FST and measured neurotransmitter levels in the hypothalamus at the end of behavioral testing. Once again, we found significant differences between the knockout lines and wild type mice as well as significant effects of BaP exposure. In our first experiment, BaP exposure increased corticosterone levels in wild type mice compared to corn oil controls, but decreased them in BaP-exposed *Cyp1a1(−/−)* compared to corn oil-treated knockouts. The pattern of a reduced stress response and reduced immobility in FST are consistent for *Cyp1a1(−/−)* knockout mice, but cannot explain the FST results for wild type mice. We found a strikingly different pattern in the *Cyp1b1(−/−)* knockout mice. Corticosterone levels were higher in both corn oil- and BaP-exposed knockouts compared with wild type mice, even though the knockouts spent less time immobile in the FST. This is the opposite pattern from the *Cyp1a1(−/−)* knockouts and suggests a different mechanism or a differential response to a stressful stimulus.

Serotonin (5-HT) is a major monoamine neurotransmitter associated with changes in mood and behavior (Tejeda-Martinez et al. 2024) that is metabolized to 5-HIAA. We found decreased serotonin turnover in *Cyp1a1(−/−)* knockout mice (Fig. 5A) and in mice exposed to BaP during early brain development (Fig. 5B) evidenced by significantly decreased levels of 5-HIAA in the hypothalamus. In contrast, *Cyp1b1(−/−)* knockout mice had lower levels of serotonin in the hypothalamus (Fig. 5D). It is now recognized that CYP enzymes have a role in tryptophan metabolism and that the presence of tryptophan metabolites can affect the level of CYP1enzymes in humans and other animals (Hadduch et al. 2023). Given the importance of tryptophan metabolism in multiple neurological disorders (Marx et al. 2020) and the emerging understanding of the tryptophan-kynurenine pathway in the gut-brain axis (Gheorghe et al. 2019, Vazquez-Medina et al. 2024), it will be important to conduct further studies on the role of CYP1 enzymes in both the gut and in the brain to fully understand their roles and the impact of mutations that alter their metabolic capacities.

Dopamine’s role in the hypothalamus is complex and intersects with numerous other neuroendocrine and neurotransmitter pathways (reviewed in Ugromov 2024). Dopaminergic signaling is central to normal circadian rhythms (Mesgar et al. 2023), and stress can impair normal dopaminergic functioning in both rodents (Ercan et al. 2023) and fish (Gesto et al. 2008). We found decreased dopamine and its metabolite DOPAC in BaP-exposed wild type male mice (Tables 1-2) and significantly decreased DOPAC in *Cyp1b1(−/−)* knockout mice (Fig. 5C). Further work is needed to determine whether these changes are related to dopamine synthesis or biodegradation; however, previous studies have implicated aryl hydrocarbon receptor agonists with dopaminergic dysfunction during brain development (Tanida et al. 2014). The mechanism appears to be direct modulation of dopamine synthesis based on the studies of Akahoshi et al. (2009) who found dioxin (TCDD) increased expression of tyrosine hydroxylase mediated by binding of the aryl hydrocarbon receptor. Overproduction of dopamine can increase oxidative stress in neurons. Therefore, it’s plausible that developmental exposure to the AHR agonist BaP can act similarly in upregulating tyrosine hydroxylase ultimately leading to oxidative stress and neurotoxicity.

Although new and novel mechanisms may play a role in differential responses to developmental BaP exposure, differential toxicokinetics must never be ignored. Adult mice with variation in the aryl hydrocarbon receptor and CYP1 enzymes have clear and striking differences in the metabolism and clearance of BaP (Uno et al. 2004, 2006). We found similar differences when measuring BaP and metabolite levels in wild type and *Cyp1* knockout dams and their pups (Feltner et al. 2023). Given recent human studies demonstrating that CYP1A1 polymorphisms can increase the concentration of PAHs crossing the placenta into cord blood (Dong et al. 2018), it seems clear that the results from animal studies have translational value for humans.

## 5. Conclusion

Exposure to BaP and other PAHs remains widespread through air, food and soil (Bukowska et al. 2022) and will likely increase in coming years as climate change leads to more intense and frequent wildfires (Ghetu et al. 2022, Jain et al. 2024). We found that developmental BaP exposure altered corticosterone levels and neurotransmitter levels in the hypothalamus, but that genotype and sex were also key factors. Mutations in both CYP1A1 (Frikha et al. 2024) and CYP1B1 (Shahid et al. 2022) are well known in the human population; therefore, our work has potential translational value by identifying individuals at higher risk of neurobehavioral impairments following BaP exposure. Our multiple findings of behavioral changes in both knockout lines of mice regardless of treatment are also likely to lead to new insights into CYP1 enzymes and their roles in neuroendocrine homeostasis and neurotransmitter metabolism.

## 6. CRediT author statement

**Jade Perry:** Conceptualization, Methodology, Investigation, Funding acquisition. **Taylor Easybuck:** Writing-Original draft, Data curation, Investigation, Visualization. **Mackenzie Feltner**: Investigation, Data curation, Supervision. **Emma G. Foster:** Investigation, Data curation, Supervision. **Mickayla Kowalski:** Investigation, Funding acquisition. **Amanda Honaker**: Investigation **Angela Kyntchev**: Conceptualization, Investigation, **Katelyn Clough:** Conceptualization, Investigation, Data curation, **Allie Easton**: Investigation. **Kalyani Abbaraju**: Investigation. **Joseph Ashley**: Investigation, **Kevin Berling**: Investigation**. India Davis**: Investigation. **Duong Pham**: Investigation. **Annika White**: Investigation, Visualization. **Kayla Wypasek**: Investigation. **Christine Perdan Curran:** Project administration, Funding acquisition, Analysis, Supervision, Writing-Original draft and Revision.

## 7. Data Availability Statement

All data and detailed protocols are freely available upon request to the corresponding author.

## 8. Acknowledgments

We acknowledge support from multiple funding sources, including NIH grants R15ES030541, R15ES020053, P20GM103436, and NSF RSF-034-07, Society of Toxicology internship grants, and the following Northern Kentucky University sources: College of Arts and Sciences Collaborative Faculty-Student Project Awards, Faculty Development Project Grants, Center for Integrative Natural Sciences and Mathematics (CINSAM) UR-STEM Fellowships, Greaves and Herrmann Fellowships, and Student Undergraduate Research and Creative Activities awards. We thank Dr. Daniel W. Nebert of the University of Cincinnati Medical Center for the generous donation of *Cyp1(−/−)* knockout mice.

## References

Akahoshi, E., Yoshimura, S., Uruno, S., Ishihara-Sugano, M., 2009. Effect of dioxins on regulation of tyrosine hydroxylase gene expression by aryl hydrocarbon receptor: a neurotoxicology study. Environ Health 8, 24. 10.1186/1476-069X-8-24

Aparna, S., Patri, M., 2021. Benzo[a]pyrene exposure and overcrowding stress impacts anxiety-like behavior and impairs learning and memory in adult zebrafish,. Environmental Toxicology 36, 352–361. 10.1002/tox.23041

ATSDR (Agency for Toxic Substances and Disease Registry) 1995. Toxicological profile for polycyclic aromatic hydrocarbons. Atlanta, GA: U.S. Department of Health and Human Services, Public Health Service.

ATSDR, 2022. Substance Priority List | ATSDR [WWW Document]. URL https://www.atsdr.cdc.gov/spl/index.html (accessed 8.1.23).

Bouayed, J., Bohn, T., Tybl, E., Kiemer, A.K., Soulimani, R., 2012. Benzo[α]pyrene-Induced Anti-Depressive-like Behaviour in Adult Female Mice: Role of Monoaminergic Systems. Basic & Clinical Pharmacology & Toxicology 110, 544–550. 10.1111/j.1742-7843.2011.00853.x

Brown, J., Villalona, Y., Weimer, J., Ludwig, C.P., Hays, B.T., Massie, L., Marczinski, C.A., Curran, C.P., 2020. Supplemental taurine during adolescence and early adulthood has sex-specific effects on cognition, behavior and neurotransmitter levels in C57BL/6J mice dependent on exposure window. Neurotoxicol Teratol 79, 106883. 10.1016/j.ntt.2020.106883

Bukowska, B., Mokra, K., Michałowicz, J., 2022. Benzo[a]pyrene-Environmental Occurrence, Human Exposure, and Mechanisms of Toxicity. Int J Mol Sci 23, 6348. 10.3390/ijms23116348

Buters, J.T., Sakai, S., Richter, T., Pineau, T., Alexander, D.L., Savas, U., Doehmer, J., Ward, J.M., Jefcoate, C.R., Gonzalez, F.J., 1999. Cytochrome P450 CYP1B1 determines susceptibility to 7, 12-dimethylbenz[a]anthracene-induced lymphomas. Proc Natl Acad Sci U S A 96, 1977–1982. 10.1073/pnas.96.5.1977

Colettis, N., Higgs, J., Wasowski, C., Knez, D., Gobec, S., Pastore, V., Marder, M., 2024. 3,3-Dibromoflavanone, a synthetic flavonoid derivative for pain management with antidepressant-like effects and fewer side effects than those of morphine in mice. Chem Biol Interact 402, 111189. 10.1016/j.cbi.2024.111189

Dalton, T.P., Dieter, M.Z., Matlib, R.S., Childs, N.L., Shertzer, H.G., Genter, M.B., Nebert, D.W., 2000. Targeted knockout of Cyp1a1 gene does not alter hepatic constitutive expression of other genes in the mouse [Ah] battery. Biochem Biophys Res Commun 267, 184–189. 10.1006/bbrc.1999.1913

Das, S.K., Patri, M., 2017. Neuropeptide Y expression confers benzo[a]pyrene induced anxiolytic like behavioral response during early adolescence period of male Wistar rats. Neuropeptides 61, 23–30. 10.1016/j.npep.2016.07.001

Dong, X., Wang, Q., Peng, J., Wu, M., Pan, B., Xing, B., 2018. Transfer of polycyclic aromatic hydrocarbons from mother to fetus in relation to pregnancy complications. Science of The Total Environment 636, 61–68. 10.1016/j.scitotenv.2018.04.274

Ercan, Z., Bulmus, O., Kacar, E., Serhatlioglu, I., Zorlu, G., Kelestimur, H., 2023. Treadmill Exercise Improves Behavioral and Neurobiological Alterations in Restraint-Stressed Rats. J Mol Neurosci 73, 831–842. 10.1007/s12031-023-02159-2

Feltner, M., Hare, P.M., Good, A., Foster, E.G., Clough, K., Perry, J., Honaker, A., Kyntchev, A., Kowalski, M., Curran, C.P., 2023. Differential Susceptibility to Benzo[a]pyrene Exposure during Gestation and Lactation in Mice with Genetic Variations in the Aryl Hydrocarbon Receptor and Cyp1 Genes. Toxics 11, 778. 10.3390/toxics11090778

Frikha, I., Frikha, R., Medhaffer, M., Charfi, H., Turki, F., Elloumi, M., 2024. Impact of CYP1A1 variants on the risk of acute lymphoblastic leukemia: evidence from an updated meta-analysis. Blood Res 59, 9. 10.1007/s44313-024-00007-9

Gesto, M., Soengas, J.L., Míguez, J.M., 2008. Acute and prolonged stress responses of brain monoaminergic activity and plasma cortisol levels in rainbow trout are modified by PAHs (naphthalene, β-naphthoflavone and benzo(a)pyrene) treatment. Aquatic Toxicology 86, 341–351. 10.1016/j.aquatox.2007.11.014

Gheorghe, C.E., Martin, J.A., Manriquez, F.V., Dinan, T.G., Cryan, J.F., Clarke, G., 2019. Focus on the essentials: tryptophan metabolism and the microbiome-gut-brain axis. Curr Opin Pharmacol 48, 137–145. 10.1016/j.coph.2019.08.004

Ghetu, C.C., Rohlman, D., Smith, B.W., Scott, R.P., Adams, K.A., Hoffman, P.D., Anderson, K.A., 2022. Wildfire Impact on Indoor and Outdoor PAH Air Quality. Environ Sci Technol 56, 10042–10052. 10.1021/acs.est.2c00619

Grova, N., Schroeder, H., Farinelle, S., Prodhomme, E., Valley, A., Muller, C.P., 2008. Sub-acute administration of benzo[*a*]pyrene (B[*a*]P) reduces anxiety-related behaviour in adult mice and modulates regional expression of *N*-methyl-d-aspartate (NMDA) receptors genes in relevant brain regions. Chemosphere, Halogenated Persistent Organic Pollutants Dioxin 2005 73, S295–S302. 10.1016/j.chemosphere.2007.12.037

Grova, N., Valley, A., Turner, J.D., Morel, A., Muller, C.P., Schroeder, H., 2007. Modulation of behavior and NMDA-R1 gene mRNA expression in adult female mice after sub-acute administration of benzo(a)pyrene. Neurotoxicology 28, 630–636. 10.1016/j.neuro.2007.01.010

Gutierrez, A., Williams, M.T., Vorhees, C.V., 2018. A single high dose of methamphetamine reduces monoamines and impairs egocentric and allocentric learning and memory in adult male rats. Neurotox Res 33, 671–680. 10.1007/s12640-018-9871-9

Haduch, A., Bromek, E., Kuban, W., Daniel, W.A., 2023. The Engagement of Cytochrome P450 Enzymes in Tryptophan Metabolism. Metabolites 13, 629. 10.3390/metabo13050629

Hamilton, T.J., Krook, J., Szaszkiewicz, J., Burggren, W., 2021. Shoaling, boldness, anxiety-like behavior and locomotion in zebrafish (Danio rerio) are altered by acute benzo[a]pyrene exposure. Sci Total Environ 774, 145702. 10.1016/j.scitotenv.2021.145702

Honaker, A., Kyntchev, A., Foster, E., Clough, K., Hawk, G., Asiedu, E., Berling, K., DeBurger, E., Feltner, M., Ferguson, V., Forrest, P.T., Jenkins, K., Massie, L., Mullaguru, J., Niang, M.D., Perry, C., Sene, Y., Towell, A., Curran, C.P., 2022. The behavioral effects of gestational and lactational benzo[a]pyrene exposure vary by sex and genotype in mice with differences at the Ahr and Cyp1a2 loci. Neurotoxicology and Teratology 89, 107056. 10.1016/j.ntt.2021.107056

Jain, P., Barber, Q.E., Taylor, S.W., Whitman, E., Castellanos Acuna, D., Boulanger, Y., Chavardès, R.D., Chen, J., Englefield, P., Flannigan, M., Girardin, M.P., Hanes, C.C., Little, J., Morrison, K., Skakun, R.S., Thompson, D.K., Wang, X., Parisien, M.-A., 2024. Drivers and Impacts of the Record-Breaking 2023 Wildfire Season in Canada. Nat Commun 15, 6764. 10.1038/s41467-024-51154-7

Kilkenny, C., Browne, W.J., Cuthill, I.C., Emerson, M., Altman, D.G., 2010. Improving bioscience research reporting: The ARRIVE guidelines for reporting animal research. J Pharmacol Pharmacother 1, 94–99. 10.4103/0976-500X.72351

Margolis, A.E., Herbstman, J.B., Davis, K.S., Thomas, V.K., Tang, D., Wang, Y., Wang, S., Perera, F.P., Peterson, B.S., Rauh, V.A., 2016. Longitudinal effects of prenatal exposure to air pollutants on self-regulatory capacities and social competence. J Child Psychol Psychiatry 57, 851–860. 10.1111/jcpp.12548

Margolis, A.E., Ramphal, B., Pagliaccio, D., Banker, S., Selmanovic, E., Thomas, L.V., Factor-Litvak, P., Perera, F., Peterson, B.S., Rundle, A., Herbstman, J.B., Goldsmith, J., Rauh, V., 2021. Prenatal exposure to air pollution is associated with childhood inhibitory control and adolescent academic achievement. Environ Res 202, 111570. 10.1016/j.envres.2021.111570

Marx, W., McGuinness, A.J., Rocks, T., Ruusunen, A., Cleminson, J., Walker, A.J., Gomes-da-Costa, S., Lane, M., Sanches, M., Diaz, A.P., Tseng, P.-T., Lin, P.-Y., Berk, M., Clarke, G., O’Neil, A., Jacka, F., Stubbs, B., Carvalho, A.F., Quevedo, J., Soares, J.C., Fernandes, B.S., 2020. The kynurenine pathway in major depressive disorder, bipolar disorder, and schizophrenia: a meta-analysis of 101 studies. Mol Psychiatry. 10.1038/s41380-020-00951-9

Mesgar, S., Eskandari, K., Karimian-Sani-Varjovi, H., Salemi-Mokri-Boukani, P., Haghparast, A., 2023. The Dopaminergic System Modulates the Electrophysiological Activity of the Suprachiasmatic Nucleus Dependent on the Circadian Cycle. Neurochem Res 48, 3420–3429. 10.1007/s11064-023-03988-8

Percie du Sert, N., Hurst, V., Ahluwalia, A., Alam, S., Avey, M.T., Baker, M., Browne, W.J., Clark, A., Cuthill, I.C., Dirnagl, U., Emerson, M., Garner, P., Holgate, S.T., Howells, D.W., Karp, N.A., Lazic, S.E., Lidster, K., MacCallum, C.J., Macleod, M., Pearl, E.J., Petersen, O.H., Rawle, F., Reynolds, P., Rooney, K., Sena, E.S., Silberberg, S.D., Steckler, T., Würbel, H., 2020. The ARRIVE guidelines 2.0: Updated guidelines for reporting animal research. Br J Pharmacol 177, 3617–3624. 10.1111/bph.15193

Perera, F.P., Tang, D., Wang, S., Vishnevetsky, J., Zhang, B., Diaz, D., Camann, D., Rauh, V., 2012. Prenatal polycyclic aromatic hydrocarbon (PAH) exposure and child behavior at age 6-7 years. Environ Health Perspect 120, 921–926. 10.1289/ehp.1104315

Perera, F.P., Wheelock, K., Wang, Y., Tang, D., Margolis, A.E., Badia, G., Cowell, W., Miller, R.L., Rauh, V., Wang, S., Herbstman, J.B., 2018. Combined effects of prenatal exposure to polycyclic aromatic hydrocarbons and material hardship on child ADHD behavior problems. Environ Res 160, 506–513. 10.1016/j.envres.2017.09.002

Shahid, M., Azfaralariff, A., Tufail, M., Hussain Khan, N., Abdulkareem Najm, A., Firasat, S., Zubair, M., Fazry, S., Law, D., 2022. Screening of high-risk deleterious missense variations in the CYP1B1 gene implicated in the pathogenesis of primary congenital glaucoma: A comprehensive in silico approach. PeerJ 10, e14132. 10.7717/peerj.14132

Tanida, T., Tasaka, K., Akahoshi, E., Ishihara-Sugano, M., Saito, M., Kawata, S., Danjo, M., Tokumoto, J., Mantani, Y., Nagahara, D., Tabuchi, Y., Yokoyama, T., Kitagawa, H., Kawata, M., Hoshi, N., 2014. Fetal exposure to 2,3,7,8-tetrachlorodibenzo-p-dioxin transactivates aryl hydrocarbon receptor-responsive element III in the tyrosine hydroxylase immunoreactive neurons of the mouse midbrain. J Appl Toxicol 34, 117–126. 10.1002/jat.2839

Tejeda-Martínez, A.R., Ramos-Molina, A.R., Brand-Rubalcava, P.A., Flores-Soto, M.E., 2024. Involvement of serotonergic receptors in depressive processes and their modulation by β-arrestins: A review. Medicine 103, e38943. 10.1097/MD.0000000000038943

Ugrumov, M.V., 2024. Hypothalamic neurons fully or partially expressing the dopaminergic phenotype: development, distribution, functioning and functional significance. A review. Front Neuroendocrinol 75, 101153. 10.1016/j.yfrne.2024.101153

Uno, S., Dalton, T.P., Derkenne, S., Curran, C.P., Miller, M.L., Shertzer, H.G., Nebert, D.W., 2004. Oral exposure to benzo[a]pyrene in the mouse: detoxication by inducible cytochrome P450 is more important than metabolic activation. Mol Pharmacol 65, 1225–1237. 10.1124/mol.65.5.1225

Uno, S., Dalton, T.P., Dragin, N., Curran, C.P., Derkenne, S., Miller, M.L., Shertzer, H.G., Gonzalez, F.J., Nebert, D.W., 2006. Oral benzo[a]pyrene in Cyp1 knockout mouse lines: CYP1A1 important in detoxication, CYP1B1 metabolism required for immune damage independent of total-body burden and clearance rate. Mol Pharmacol 69, 1103–1114. 10.1124/mol.105.021501

Vahid-Ansari, F., Newman-Tancredi, A., Fuentes-Alvarenga, A.F., Daigle, M., Albert, P.R., 2024. Rapid reorganization of serotonin projections and antidepressant response to 5-HT1A-biased agonist NLX-101 in fluoxetine-resistant cF1ko mice. Neuropharmacology 110132. 10.1016/j.neuropharm.2024.110132

Vazquez-Medina, A., Rodriguez-Trujillo, N., Ayuso-Rodriguez, K., Marini-Martinez, F., Angeli-Morales, R., Caussade-Silvestrini, G., Godoy-Vitorino, F., Chorna, N., 2024. Exploring the interplay between running exercises, microbial diversity, and tryptophan metabolism along the microbiota-gut-brain axis. Front. Microbiol. 15. 10.3389/fmicb.2024.1326584

Vishnevetsky, J., Tang, D., Chang, H.-W., Roen, E.L., Wang, Y., Rauh, V., Wang, S., Miller, R.L., Herbstman, J., Perera, F.P., 2015. Combined effects of prenatal polycyclic aromatic hydrocarbons and material hardship on child IQ. Neurotoxicol Teratol 49, 74–80. 10.1016/j.ntt.2015.04.002

Zhang, W., Tian, F., Zheng, J., Li, S., Qiang, M., 2016. Chronic administration of benzo(a)pyrene induces memory impairment and anxiety-like behavior and increases of NR2B DNA methylation. PLOS ONE 11, e0149574. 10.1371/journal.pone.0149574

Zhen, H., Zhang, F., Cheng, H., Hu, F., Jia, Y., Hou, Y., Shang, M., Yu, H., Jiang, M., 2023. Association of polycyclic aromatic hydrocarbons exposure with child neurodevelopment and adult emotional disorders: A meta-analysis study. Ecotoxicology and Environmental Safety 255, 114770. 10.1016/j.ecoenv.2023.114770

